# Implementing the reuse of public DIA proteomics datasets: from the PRIDE database to Expression Atlas

**DOI:** 10.1101/2021.06.08.447493

**Authors:** Mathias Walzer, David García-Seisdedos, Ananth Prakash, Paul Brack, Peter Crowther, Robert L. Graham, Nancy George, Suhaib Mohammed, Pablo Moreno, Irene Papathedourou, Simon J. Hubbard, Juan Antonio Vizcaíno

## Abstract

The number of mass spectrometry (MS)-based proteomics datasets in the public domain keeps increasing, particularly those generated by Data Independent Acquisition (DIA) approaches such as SWATH-MS. Unlike Data Dependent Acquisition datasets, the re-use of DIA datasets has been rather limited to date, despite its high potential, due to the technical challenges involved. We introduce a (re-)analysis pipeline for public SWATH-MS datasets which includes a combination of metadata annotation protocols, automated workflows for MS data analysis, statistical analysis, and the integration of the results into the Expression Atlas resource. Automation is orchestrated with Nextflow, using containerised open analysis software tools, rendering the pipeline readily available and reproducible. To demonstrate its utility, we reanalysed 10 public DIA datasets from the PRIDE database, comprising 1,278 SWATH-MS runs. The robustness of the analysis was evaluated, and the results compared to those obtained in the original publications. The final expression values were integrated into Expression Atlas, making SWATH-MS experiments more widely available and combining them with expression data originating from other proteomics and transcriptomics datasets.

## Introduction

The availability of mass spectrometry (MS)-based proteomics datasets continues to increase dramatically, representing a growing and increasingly useful resource for the biomedical sciences. Indeed, this growth mirrors that found in other related omics fields such as transcriptomics, where re-use, re-analysis and new comparative studies can be facilitated^1,2^. The PRIDE (PRoteomics IDEntifications) database^3^ at the European Bioinformatics Institute (EBI, https://www.ebi.ac.uk/pride/) is the largest proteomics data repository worldwide and is one of the founding members of the ProteomeXchange consortium^4^. During 2021 alone, PRIDE captured around 5,800 datasets, originating from a wide variety of species and different experimental approaches.

One of the main benefits of making data publicly available is to enable reuse and increases reproducibility, facilitating an independent assessment of the results described in the corresponding publications. This represents an auditable route to (re)trace the source of key findings as well as supporting the potential discovery of new findings as new advanced software becomes available. The growth in public domain proteomics data has indeed triggered data reuse activities^5,6^ and new applications in the field, including among others, numerous meta-analysis studies, proteogenomics applications and the use of artificial intelligence approaches such as machine-learning^7,8^. Expression Atlas (https://www.ebi.ac.uk/gxa/home) is an added-value resource at the EBI that enables easy access to integrated information about gene and protein expression across species, tissues, cells, experimental conditions and diseases^9^. The Expression Atlas (EA) ‘bulk’ database has two sections: baseline and differential, and already integrates some proteomics data. This data has been generated from reanalysis of human, mouse, rat and cell-line public datasets, all of which have been derived from Data Dependent Acquisition (DDA)^9–11^. The availability of such results in EA therefore supports the integration of proteomics with transcriptomics information.

To date, only DDA-based proteomics datasets have been reanalysed and integrated in EA. However, an increasing fraction of data deposited in public repositories comes from DIA (Data Independent Acquisition) approaches and from SWATH-MS (Sequential Window Acquisition of All Theoretical Mass Spectra) methods in particular. DIA methods differ from DDA techniques in that no narrow-window ion selection takes place in the mass spectrometer. As a result, fragment ion spectra of all the available peptide ion species in a wide window are produced. In the case of SWATH-MS, this is conducted in cycles of sequential windows of *m/z* (mass-to-charge) values^12,13^, creating permanent digital maps of the protein samples via the precursor and fragment ions of their proteolytic peptides. These digital maps can potentially be re-interrogated over time^13–16^, for instance with novel spectral libraries including additional peptides. This holds great promise for biomarker discovery and other applications, especially since DIA approaches can have distinct advantages for quantitative proteomics studies. For instance, high reproducibility between technical and biological replicates^17^, and high intra-and inter-laboratory reproducibility^18^. As a result, SWATH-MS methods can capture a comprehensive picture of the sample measured and have established themselves as a reproducible method for large-scale protein quantification.

Despite this great promise, SWATH-MS analysis pipelines are complex and involve multiple stages and parameters. Typically, the *m/z* space per cycle is segmented into a set of acquisition windows, within which the analysis software attempts to detect signals from a library of expected peptides provided in a spectral library. Depending on the instrument speed, target acquisition *m/z* range and gradient length, either 32 or 64 equally sized windows are used, though this can vary from study to study, and dynamically sized windows are also often used. Similarly, the methods used to detect, score and statistically validate the underlying peptide-associated mass spectral features (i.e., co-eluting ion group signals from different transitions) can vary greatly between studies, and depend on the data analysis software used. This task is often accomplished via a targeted approach where peptides from a spectral library are searched for in the digital map using a look-up table of precursor ion *m/z* values, expected fragment ions (*m/z, c*), and a given retention time (RT) range when they elute off a column. These are usually empirically determined on the same or a similar instrumentation and settings, generated via DDA data from pooled or representative samples. At present, despite some notable exceptions (^15,19–21^), sample/study specific target spectral libraries are rarely deposited alongside the original SWATH-MS data in public repositories such as PRIDE. Additionally, the availability of the spectral libraries in different (custom) formats make them often difficult to adapt for reanalysis. In 2014, a pan-human target library comprising 10,000 proteins was published^15^, covering 50.9% of the annotated human proteins in UniProtKB/Swiss-Prot. These targets can be used more generally as their normalised RTs are reported and additional SWATH-MS studies can be normalised to the same scale, usually by the inclusion of known iRT peptides^22^. However, the use of a generic target library requires an appropriate statistical control of peptide and protein error rates^23^.

This robustness of the overall SWATH-MS approach has been demonstrated several times, notably in a benchmarking comparison of DIA software tools which observed very similar qualitative and quantitative results^24^. Although such studies show that DIA methods are a powerful approach for reproducible protein quantification, as highlighted above, their reuse is at present very limited. This stems from the practicalities of deploying a data analysis pipeline, due to either the large amounts of data to be processed, the huge computational efforts needed, the intricate software settings necessary, or the uncertainty about whether the data from a particular study is adequate for reanalysis in a different research setting. In this study, we provide a robust open reanalysis pipeline for DIA data and demonstrate its utility to confidently reanalyse ten public human DIA datasets obtained from PRIDE. Furthermore, the resulting quantified proteins are subject to further statistical analysis and the resulting protein expression data is integrated into Expression Atlas for widely available access. To the best of our knowledge, this is the first time that public DIA data has been systematically reanalysed and integrated with gene expression information. We demonstrate, that with well-formed experimental design annotation and (known) iRT peptide spiked data, it is possible to reanalyse public DIA data in a generic context, making it available to the community at https://github.com/PRIDE-reanalysis/DIA-reanalysis.

## Results

### Open analysis pipeline for DIA data

The reanalysis pipeline represents a harmonised combination of metadata annotation protocols, open proteomics software tools for SWATH-MS data in computational workflows orchestrated by Nextflow^25^ which can be run in multiple compute environments, and data integration procedures. Figure 1 illustrates the distinct parts of the overall pipeline process. The pipeline integrates all the necessary steps in SWATH-MS data analysis ranging from data curation, raw data conversion, through to statistical analysis and data integration procedures for submission into EA. Software for the different steps of each part of the pipeline are fully containerised (Table 3), making a version update to individual tools more straightforward. A new container including an updated version of a given tool can be provided through the same parameter file used to change parameter settings e.g., a FDR (False Discovery Rate) threshold for peak group detection.

**Figure 1.**
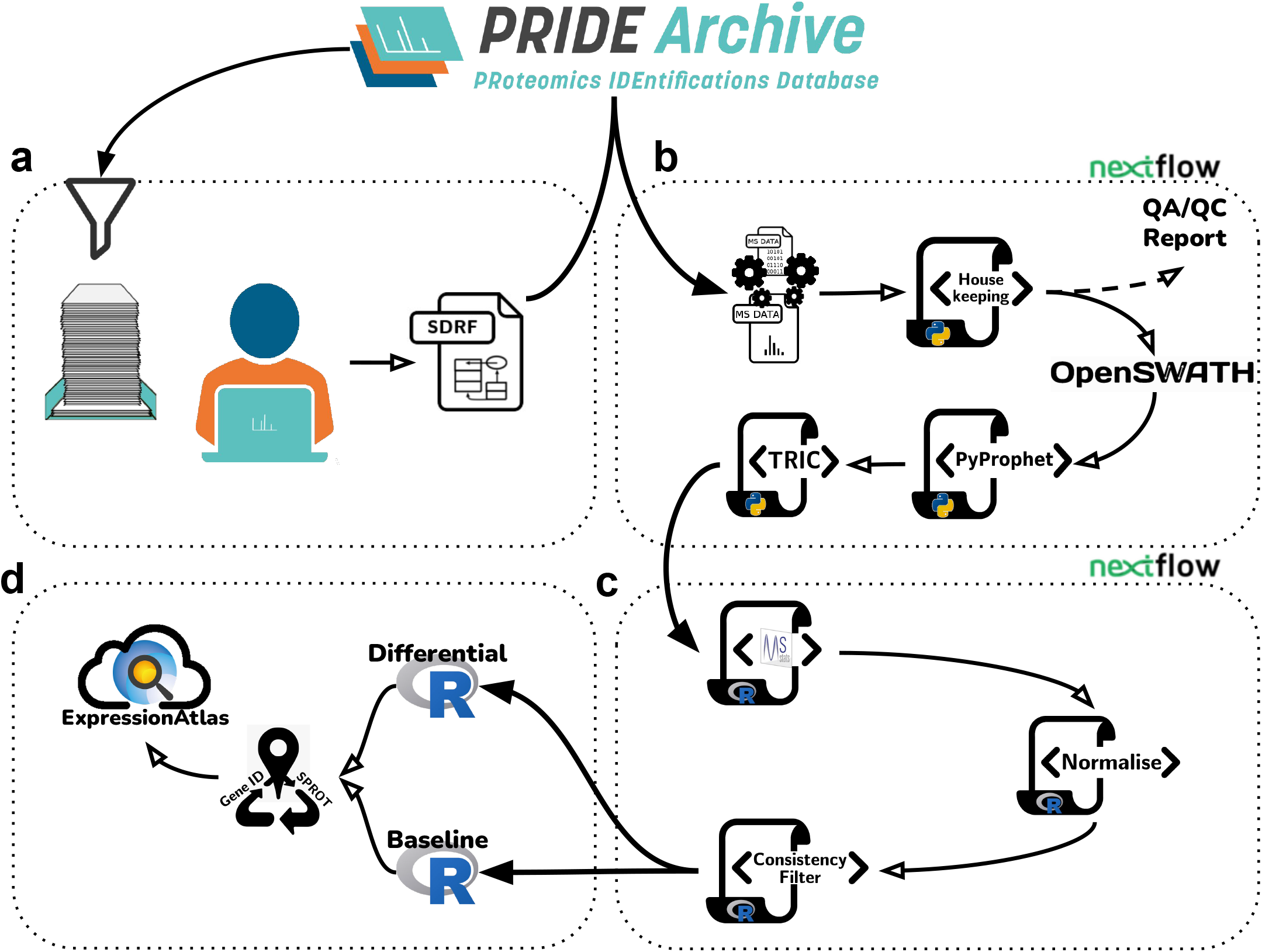
Graphical representation of the DIA data reanalysis pipeline, consisting of 4 parts. a) Data curation: Metadata annotation protocols and dataset acquisition. b) SWATH-MS data analysis: Nextflow workflow including steps ranging from data conversion, SWATH-MS window management, data quality assessment and control (QA/QC), OpenSWATH analysis, FDR calculation to measurement alignment. c) Statistical analysis: Nextflow workflow for MSstats analysis, normalisation and result filtering. d) Data integration: Data preparation, accession mapping and integration into Expression Atlas.

To add flexibility, the pipeline was split into four parts. The first (Figure 1a), includes dataset curation and acquisition, and the second (Figure 1b) contains the upstream analysis of the SWATH-MS raw data, automated with a Nextflow workflow. The third (Figure 1c), provides downstream statistical analysis, and is also automated with Nextflow, and a fourth part (Figure 1d) contains an internal workflow for the integration of the results into Expression Atlas. The split has the advantage that the analysts can inspect intermediate results before initiating the following part of the pipeline. For example, the second part contains an extra step to produce Quality Control/Assurance (QC/QA) records for each MS run, which could be inspected prior to initiating subsequent steps of the pipeline. This was particularly helpful in assessing potentially incorrect iRT peptide detection in a newly reanalysed dataset. An example for such assessment with the resulting exclusion from further analysis can be found in the Supplementary Material (Supplementary Figure S1). Optional entry points for the SWATH-MS raw data analysis workflow were added for datasets already analysed with OpenSWATH and for pre-processed spectral libraries (Supplementary Figure S2). Further details on the pipeline methods and availability can be found in the ‘Methods’ and ‘Code availability’ sections, respectively.

### Reanalysis of public DIA datasets

Ten PRIDE public datasets, amounting to 1,278 individual SWATH-MS runs (Table 1), were reanalysed as explained in the ’Methods’ section, using the EBI High Performance Computing (HPC) infrastructure. The CAL (Combined Assay Library) spectral library (SWATHAtlas accession number SAL00031) was used for all except the ’Plasma’ dataset (for which a plasma-targeted library was preferred, see ’Methods’ section).

**Table 1.**
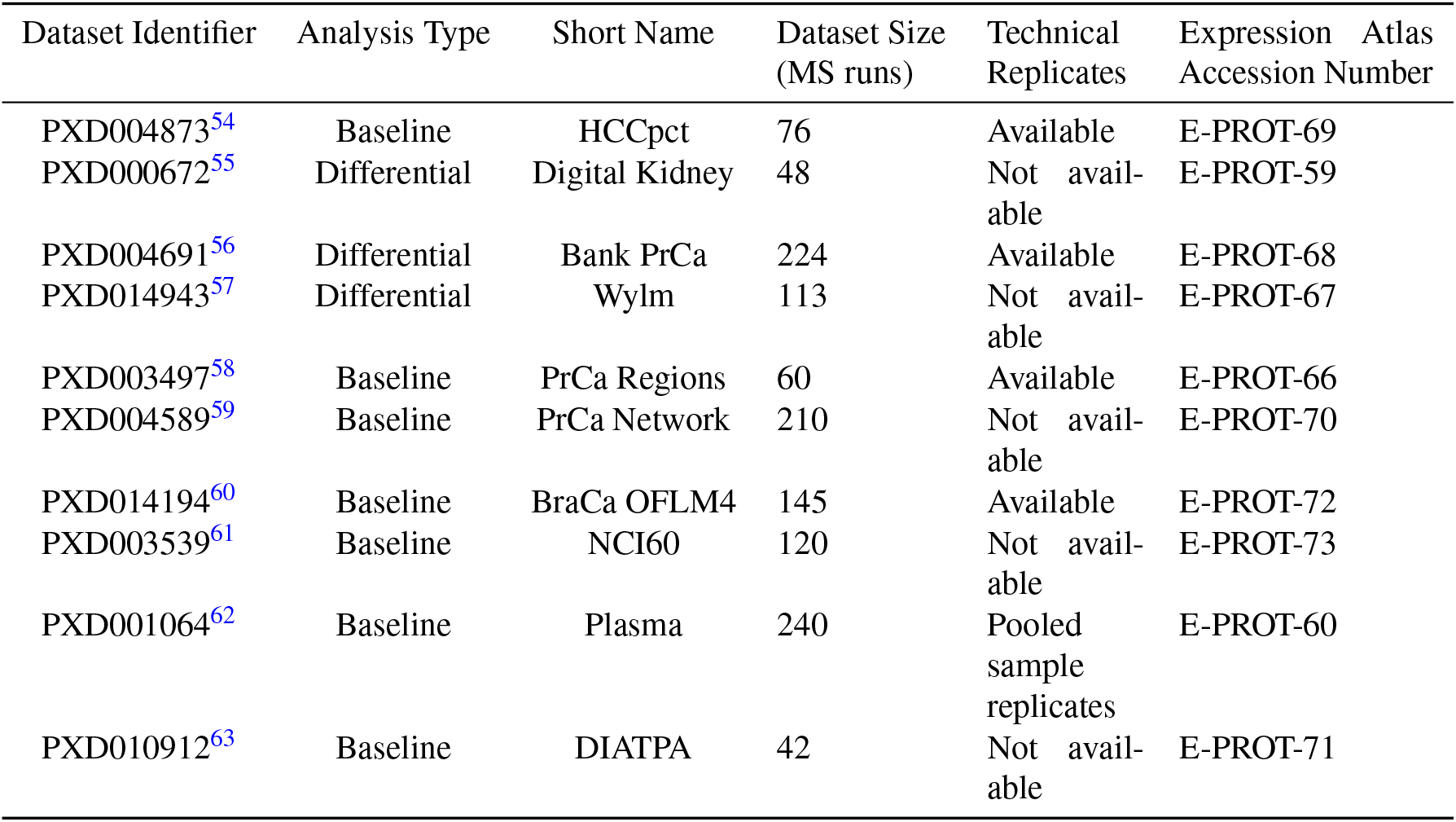
Main characteristics of the selected public DIA datasets for data reanalysis. Further details can be found in the ‘Data availability’ section.

The accumulated processing time for all datasets was around 19,000 CPU core hours, which amounted to approximately 2,200 CPU core hours (equivalent to the time that computation would take on a single core) per dataset, or 15 CPU core hours per SWATH-MS run, on average.

We first compared the raw numbers of inferred proteins across three distinct levels of protein FDR: 5%, 1% and 0.1%, filtering on the q-values from the pyProphet FDR estimates for global protein level. Overall, the numbers of detected proteins were generally in the expected range for an analysis performed with mammalian cells/tissues, using the (generic) CAL target spectral library and for protein FDR levels of 1% (Figure 2a). At a 1% cut-off for example, just under 3,000 quantified proteins were reported for PXD003497 (with 2,754) and for PXD004691 (with 2,872). Both datasets are prostate cancer sample datasets. On the higher side, PXD004589 (another prostate cancer sample dataset) yielded 3,703 detected proteins, PXD004873 (hepatocellular cancer) 3,530 proteins, and PXD010912 (human liver S9 fractions) 4,224 detected proteins. There were two datasets where fewer proteins were detected: 2,239 in PXD014194, a breast cancer dataset containing only tumour tissue measurements, and as expected, a much lower number was found in the plasma dataset PXD001064 (207 proteins). Additionally, two datasets showed many more detected proteins than the others: 5,946 in PXD014943, a lymphoma sample dataset, and 7,097 in PXD003539, the NCI-60 cell-line dataset containing samples that contributed to the design of the CAL. The FDR threshold of 1% on a global level appeared to be a good trade-off. Results for stricter and more generous cut-offs are shown in Figure 2A for comparison. We note that use of a stricter filter at 0.1% FDR proved impractical, as for the plasma dataset PXD001064 the analysis failed to complete successfully.

**Figure 2.**
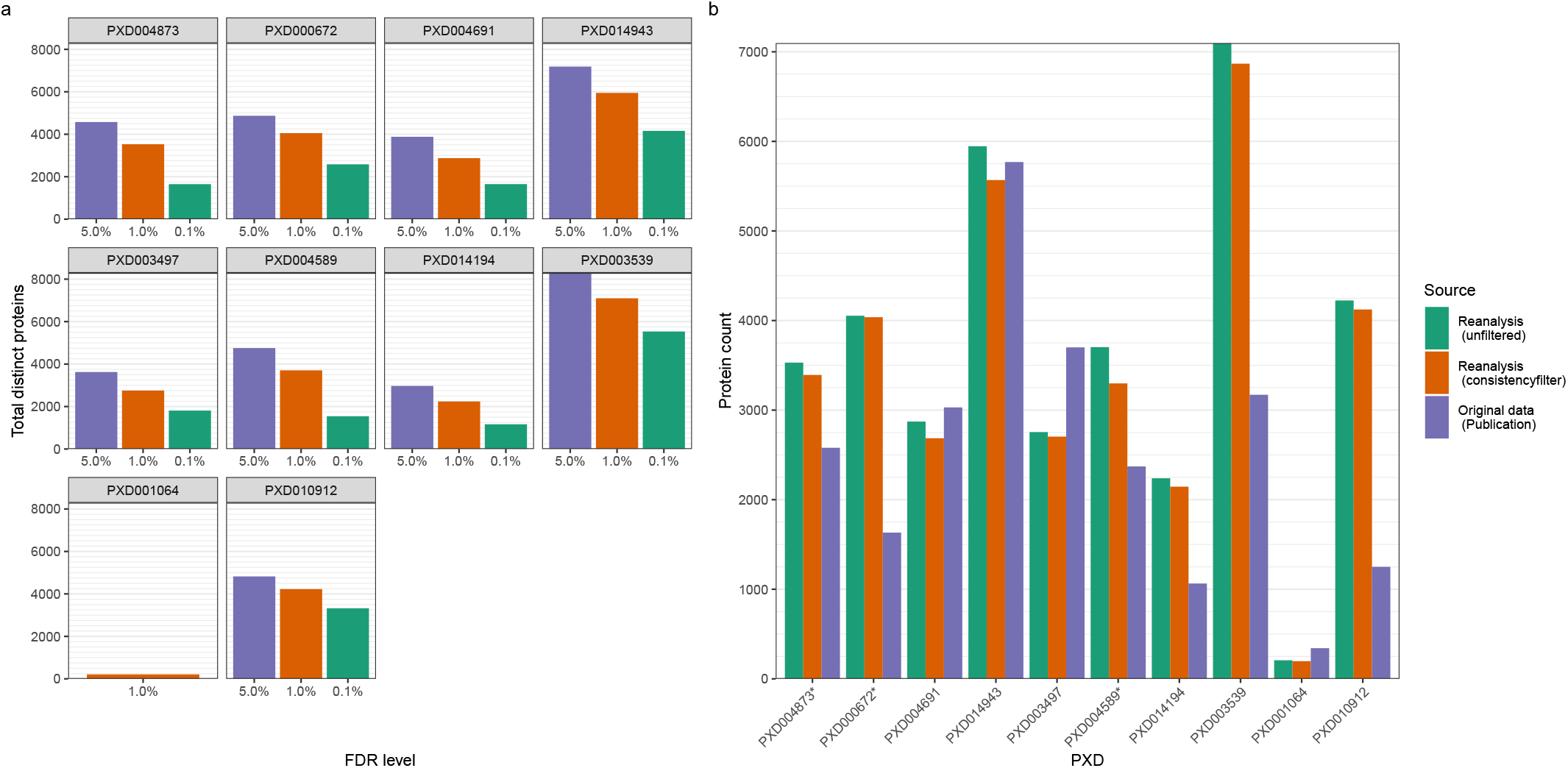
a) Number of detected proteins per dataset at different FDR levels in the data reanalysis. a) Protein detection results after 1% protein FDR threshold filtering. Original data refers to the respective publication’s mentioned protein numbers, reported at 1% protein FDR unless indicated otherwise. Reanalysis numbers are provided unfiltered and with the consistency filter applied (at least 50% of all protein’s peptide fragment targets have to be detected within a study group).*Proteins coming from datasets PXD000672 and PXD004873 were reported in the original publication at a 0.1% protein level FDR only. In the case of dataset PXD004589 at 0.1% peptide level FDR was reported.

To check the broad reliability of the open reanalysis pipeline combined with a generic spectral library, we compared our results with the originally published ones. We further implemented a consistency filter to remove unreliable protein identifications, in common with best practice in many of the published studies. Proteins or protein groups with more than 50% of target features missing within the MS runs across a study group were removed. We note here that an informed and consistent comparison between the original (as published) and reanalysed protein numbers across all studies is essentially impossible, since among other reasons, the original reported studies used slightly different consistency filter approaches or FDR thresholds. On one hand, for 6 of the 10 studies considered, after reanalysis, the number of proteins was higher than those originally reported. In 4 of them, the numbers were markedly higher (Figure 2b). This is likely due to the protein numbers for dataset PXD000672 and PXD004873 being originally reported at a 0.1% protein level FDR, at a 0.1% peptide level FDR for PXD004589. Additionally, PXD014149 was originally analysed with a target library of 3,284 proteins. In the case of PXD010912 we assume fewer proteins were present in the original spectral library (not reported in the original publication) as well, as the authors reported that only 580 proteins were quantified via DDA. The Supplementary Material to the original publication of PXD003539 offers 6,556 protein groups with missing values quantified, a much closer value to the results of the reanalysis.

On the other hand, the protein numbers of 4 studies were lower than those originally published: around 300 proteins less in all cases with the exception of dataset PXD003497, where a difference of just under 1,000 proteins was found. Interestingly, this last study was originally analysed with CAL, albeit customised with additional DDA measurements, obtaining 6,800 proteins. However, the authors subsequently reduced their set of proteins to 3,700 applying a more stringent consistency filter (at least two concordant proteotypic peptides^26^), which is more in line with our reanalysis.

All protein expression results are available via Expression Atlas (for URLs see Table 1). Source numbers are available in Supplementary Table S3. Intermediate results are available through the pipeline repository (see Supplementary Table S4).

### Baseline expression analysis - Technical validation

To validate the internal consistency of the reanalysed results, we calculated the coefficient of variation (CV) from the quantitative values within the respective study groups’ technical replicates and in total, comparing them to the original CVs, where available. Three of the original datasets included technical replicates and in the corresponding publication the authors reported protein intensities at a MS-run level, which are necessary to calculate the CV (Table 2). One publication reported only sample averages, and one included only pooled-sample replicates, preventing a more comprehensive CV comparison. The overall median CV was generally below 21% and the CV values obtained from the reanalyses were similar to the originally published CVs.

**Table 2.**
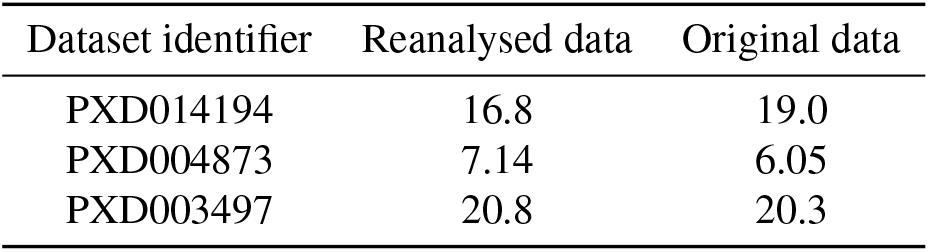
Median CV values for technical replicates, for the reanalysed results and originally published data, respectively.

The group-wise comparison showed a similar picture to the overall median CV comparison (Figure 3), although remaining in a very similar range, and were in one instance slightly smaller (Figure 3e). Again, this demonstrates the consistency of the reanalysis pipeline to reproduce the global characteristics of measurement variance observed in the original studies. To further compare the results of the reanalyses with the originally published results, we analysed the MS run-wise correlation of previously published protein abundances versus the equivalent results of the reanalyses for the technical replicates (Figure 4). The reanalysis results showed a high correlation with the original studies, though markedly closer to the regression line at a higher abundance and a wider distribution towards the lower intensities – as it might be expected. The Pearson product-moment correlation coefficients (R) ranged between 0.9 and 0.98 with a p-value ≤ 0.001, demonstrating a high concordance between the original and the reanalysed protein quantitative data overall, with matched technical reproducibility.

**Figure 3.**
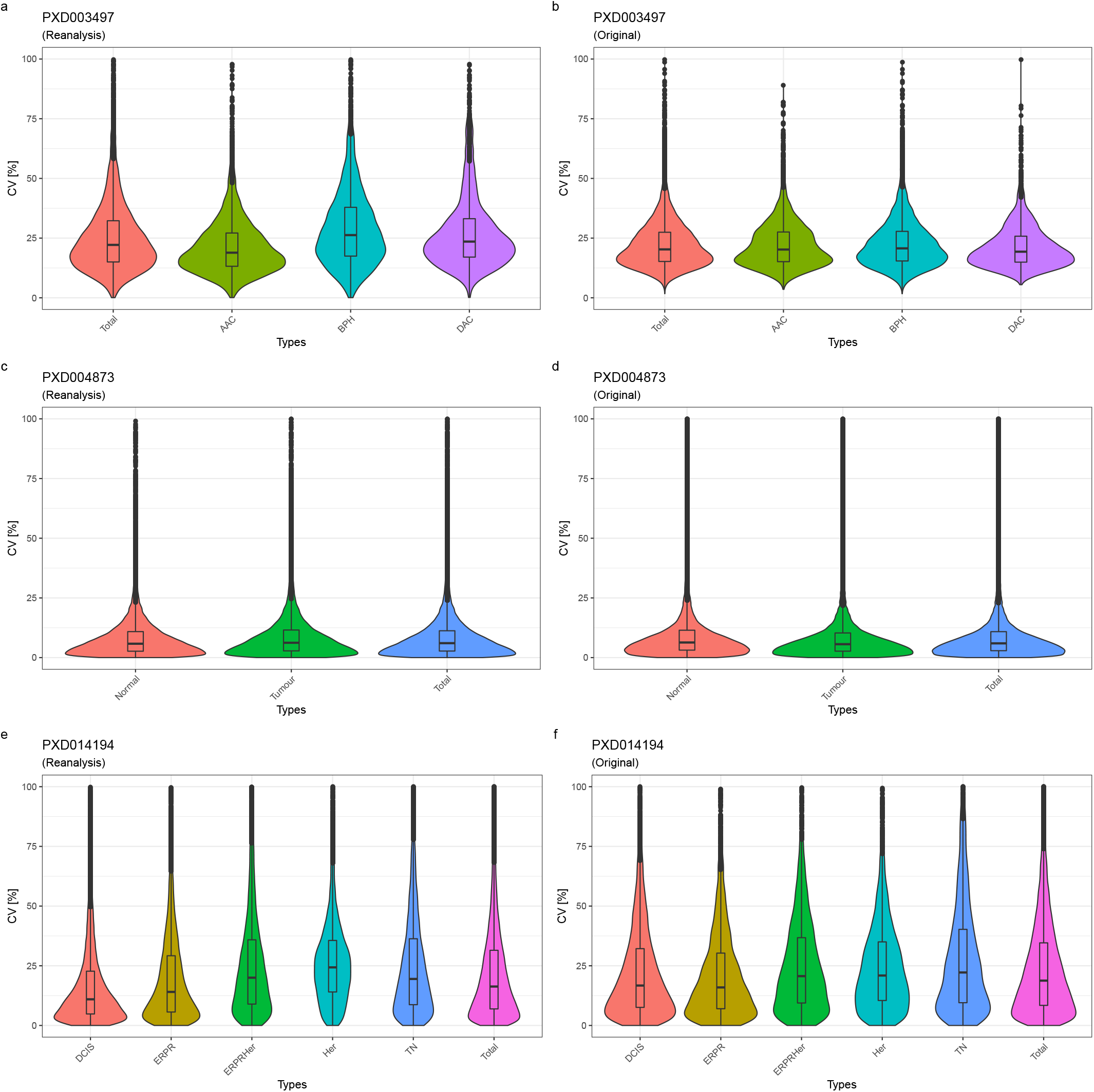
Violin-plots showing the results of the group-wise CV comparisons: a) PXD003497 reanalysis; b) PXD003497 original data; c) PXD004873 reanalysis; d) PXD004873 original data; e) PXD014194 reanalysis; f) PXD014194 original data. As it can be seen from the similar size and shapes of the violin-plots, the CVs across the datasets are largely concordant.

**Figure 4.**
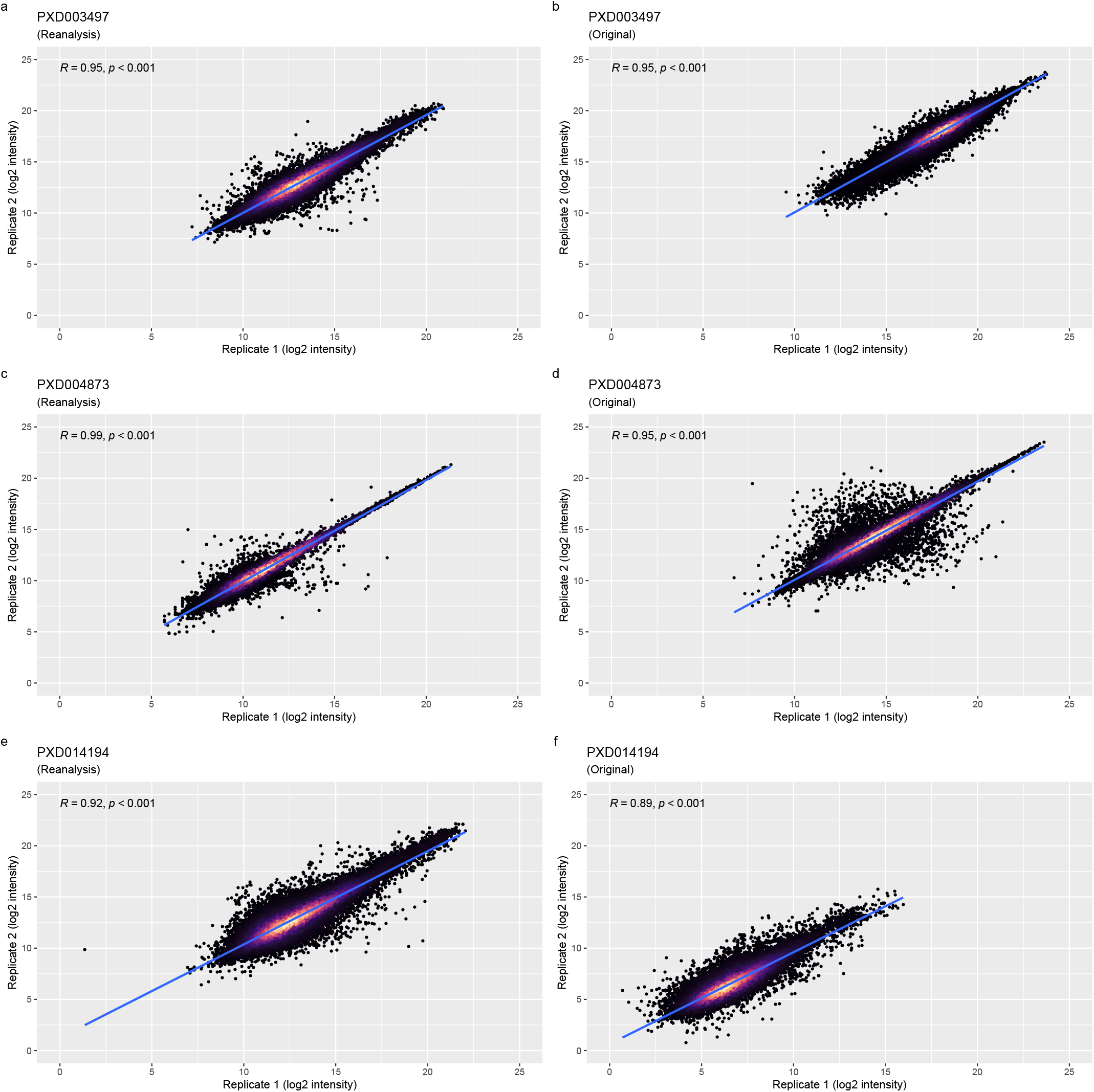
Correlation analysis of reported log2 protein intensities from technical replicate pairs: a) PXD003497 reanalysis; b) PXD003497 original data; c) PXD004873 reanalysis; d) PXD004873 original data; e) PXD014194 reanalysis; f) PXD014194 original data. The first items of pairs are on the x-axis and second items are on the y-axis. Each point represents a protein. The point density is indicated by the colour gradient, with black showing the lowest density. The higher the density the lighter the colour becomes.

**Figure 5.**
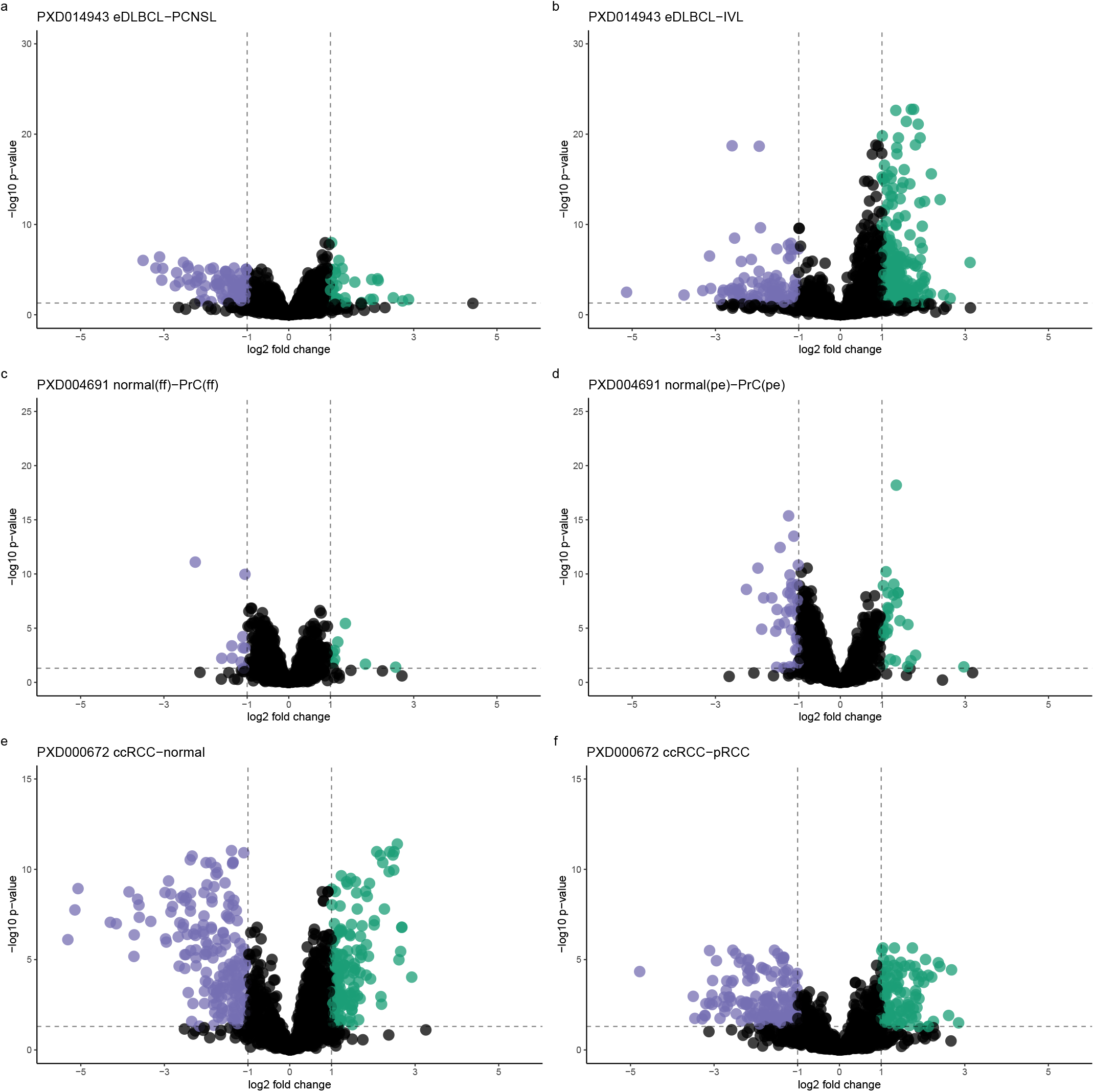
Volcano plots corresponding to the differential expression analysis for dataset PXD014943: a) extranodal diffuse large B-cell lymphoma (eDLBCL) versus primary central nervous system lymphoma (PCNSL); b) intravascular lymphoma (IVL) versus eDLBCL. For dataset PXD004691: c) normal tissue (fresh frozen) versus PrC (fresh frozen); d) normal tissue (paraffin embedded) versus tumour tissue (paraffin embedded). For dataset PXD000672: e) benign tissue samples versus clear cell RCC; f) clear cell RCC versus paillary RCC. The FC compared are represented by points on the plot. Significant FC proteins are colour indicated, dashed lines indicate the fold-change cutoff of 2 and the (adjusted) p-value cutoff at 0.05.

However, when comparing the correlation of the reanalysis results with the original results on a per-MS run pairing (reanalysis versus original results) basis, the correlations between matched protein expression values were less pronounced (R values were between 0.52 and 0.84, see Supplementary Figure S3). This is consistent with the challenges in matching the exact protein identification parameters and highlights the inherent difficulties in implementing DIA reanalyses in general.

### Differential Expression Analysis

Additionally, we also performed a downstream differential expression analysis for the three datasets that reported such values in the original publications: datasets PXD000672, PXD004691 and PXD014943. The programming language R^27^ and MSStats^28^ were used for the differential statistical analysis (see details in the ’Methods’ section). For consistency, we used the default MSstats settings (except for the use of median normalisation and the ’top3’ protein inference method). Regardless, substantial differences were found when compared with the original studies, though the number of differentially expressed proteins were in the same general order.

We calculated the correlation between the originally reported fold-changes and our reanalysis fold-changes (see Supplementary Figure S4). There was a substantial overlap of quantified protein expression values between the originally reported results and the reanalysed ones, ranging from 2,992 proteins (out of 4,572 protein fold-changes) in the reanalysis of the PXD014943 eDLBCL-PCNSL contrast, to 863 proteins (out of 1,868 protein fold-changes) in the reanalysis of the PXD000672 benign-ccRCC contrast. The higher ratio of overlap in PXD014943 also showed the highest correlation (R value at 0.84), whereas the lowest also shows the smallest correlation R value of 0.52. This illustrates that the protein signals detected were generally in the same order of magnitude yet showing a varying amount of overlap and correlation. It also hinted at an increased number of protein signals of lower intensity, that differed between the analyses to a greater extent, being picked up as the number of quantified proteins increase, and in turn reducing the overall concordance. Accordingly, when drawing a cut-off for adjusted p-values (*<* 0.05) and fold-changes (*>* 2), the overlap between the original and the reanalysed results was reduced further. For the PXD014943 eDLBCL-PCNSL contrast, we found 118 proteins to be significantly differentially expressed overlapping in 21 proteins with the original study’s 97 proteins significantly expressed (see Supplementary Table S1). The overlap of proteins of significance between original results and the reanalysed results was more pronounced in the PXD000672 benign-ccRCC contrast, with 59 of the 262 proteins from the reanalysis matching with the originally reported 613 proteins (see Supplementary Table S1). We also encountered an instance of no overlap in PXD004691 normal(ff)-PrC(ff) contrast from the 3 proteins of significance originally reported and the 8 proteins detected from the reanalysis. The corresponding protein expression values were exported into Expression Atlas as explained in the ’Methods’ section. See details about the corresponding datasets in Table 1. Of note, the analysis with the MSstats default protein inference setting of ’all’ (instead of ’top3’) resulted in a far fewer number of significantly differentially expressed proteins. Here, the intensity correlations between originally reported protein abundance values and the reanalyses (Supplementary Figure S4) revealed an unexpected lack of correspondence.

## Discussion

The popularity of DIA approaches in the proteomics field is increasing. However, to date, data reuse of public DIA datasets has been limited to mainly benchmarking studies (for example [^29,30^]). This is somewhat surprising given one of the principal advantages of SWATH-MS is to generate a permanent and comprehensive digital signature of a proteome that can be analysed again. Since this is clearly desirable, and for this to become a more common practice, best-practice for systematic reanalyses are needed to establish as well as a common reference framework. However, the complexity of a SWATH-MS study data analysis can be overwhelming. A common, ready-made pipeline lowers the entry threshold for the analysis of unseen datasets – both factors which motivated this study. Hence, in this paper, we report, to the best of our knowledge, the first systematic effort to enable the reanalysis and integration of the results of this type of studies. The implemented open reanalysis pipeline produces overall robust results and provides sufficient flexibility for users to adjust to different reanalysis scenarios.

Due to the differing experimental designs currently available, it is hard to formalise a completely generic downstream analysis workflow. However, a good compromise could be reached by designing the statistical analysis part of the pipeline using MSstats’ dataProcess and further data adjustments such as normalisation and filtering done using R. From a technical perspective, the additionally implemented entry points to the first Nextflow workflow (Figure 1b) for pre-processed data (e.g., OpenSWATH output files) proved to be advantageous, since analyses that follow the initial SWATH-MS data analysis step often needed to be tested with different parameter settings. The Nextflow configuration of iteratively repeating failed jobs with an increased memory request was also found to be advantageous. The most influential variables on memory requirements for the analysis depended on vastly differing settings between the individual studies, including liquid Chromatography (LC) gradient length, sample content, and of course the study size. HPC compute environments appear as the logical choice for data (re-)analysis efforts without clear resource requirement boundaries. However, research institutions’ collaborations with private cloud systems are becoming increasingly popular (e.g. OpenStack). SWATH-MS data analysis in the cloud, as showcased by Peters et al.^31^, can result in greatly increased compute costs, when applied to a broad basis of inputs but without the close knowledge of the dataset as the data producing lab would have - like with the datasets chosen in this study. DIA analyses are also inherently disk-intensive processes due to the size of the MS runs and the spread of analysis features throughout the data (window sequences). This, too, is an often-overlooked contributor to compute cost, especially in cloud compute environments, where storage and volume of uploaded/downloaded data are usually billed separately. The interrogation of .*wiff/*.*scan* files is, as with any other proprietary raw file formats, bound to many constraints. Inspection either needs proprietary software (e.g. Microsoft Windows), conversion into an open format which takes considerable time and disk space or is otherwise not compatible with an automated high throughput. It should be noted in this context that additional open analysis pipelines for DIA datasets have been recently developed using NextFlow^32^ and Galaxy^33^ as the workflow management systems.

During the initial selection of public datasets from PRIDE for reanalysis, we realised that a great proportion did not have (completely) paired .*wiff/*.*scan* files, rendering the unpaired samples unusable. At that point, complete pairing was made mandatory when submitting new SWATH-MS datasets to PRIDE. We hope this will help reanalysis efforts in the future, since it is essential information reflecting the experimental design as used in the original publication. Mapping raw file names in PRIDE to the samples in the original publication was done manually and it constituted one of the most time-consuming steps in this work. In the context of the activities of the Proteomics Standards Initiative, a standard file format called SDRF-Proteomics (Sample and Data Relationship Format-Proteomics) file (as part of the file format MAGE-TAB-Proteomics) has been formalised recently^34^ for capturing the experimental design in proteomics experiments^3^, and we have started working in the related tooling to facilitate the creation of these files. It is important to highlight that submission of SDRF-Proteomics files is already supported by PRIDE, although it is optional at the time of writing. The authors encourage data submitters to align and improve efforts to match metadata referenced in a publication with the underlying data and make the experimental design an explicit part of any publication. In any case, concrete and consistent name-and-group mapping tables are already of great help to any reanalysis or replication effort and increase the transparency of any original publication and therefore its value to the community. The annotation of the used iRT peptides, too, is crucial to DIA data (re-)analysis efforts. Indeed, detection of the iRT peptides is essential to their success. We consider this part of QA an important step during measurement acquisition or the reanalysis process.

We deployed the CAL target library in combination with a 1% FDR level threshold for global peptide and protein quantitation as an appropriate common setting for our reanalyses. This sometimes generated relatively low numbers of proteins quantified (when compared to the results from custom-made target libraries used in the original publications). This could be attributed to the limited number of target (peptide fragment ion) features available for some proteins in CAL. For example, the filtered CAL used for the plasma dataset reduced the number of detected proteins by over half compared to the original study. Hence, we consider the reduction of the target space for more specialised samples such as biofluids like plasma to warrant further customisation to the analysis procedure, as opposed to the *generic* application of CAL to the rest of the datasets. However, the trade-off merits between comprehensive libraries and specialised ones are beyond the scope of this initial study, and we consider the ’core’ of UniProtKB/Swiss-Prot proteins present in CAL to be appropriate for a large proportion of human tissue and cell line studies. Other strategies for future reanalysis efforts could be the use of ’library-free’ approaches^35–37^ and/or *in silico* predicted libraries^38,39^. These approaches are starting to get more popular in the field, however when this study was designed, we chose the more established open data analysis approach, which remains the most frequently used approach to date. Of course, different analysis software tools could also be used for spectral library-based approaches (e.g. Demichev et al.^30^) and we also note an extensive benchmarking study of DIA-tools was published recently^40^.

As noted, the reanalysis pipeline produced different results compared to the original published studies. However, there was a good level of agreement between replicates within each dataset, as measured by the CV values, demonstrating a robust, self-consistent quantification methodology. There was also a basic agreement between the analyses at a fold-change level, with a sufficient overlap and preservation of the general trends. However, the overlaps between the discrete sets of proteins classed as differentially expressed was often modest. We ascribe this to the large choice of parameters involved in the generation of differential expression results (independently of the analysis software used) and the serial nature of the different steps of the whole quantitative pipeline. Initial steps and choice of cut-offs (like differences in peptides detected) can become amplified to generate a large difference in later results. As shown with the (lack of) overlap in significantly different protein abundances (Supplementary Table S1), we were only partially successful in reproducing the results of the original studies.

The DIA-MS reanalyses provided a large overlap in detected peptides (Supplementary Figure S5, panels d, f) and a similarly substantial overlap in detected proteins (Supplementary Figure S5, panels g, i), despite the limited overlap in spectral library target peptides (Supplementary Figure S5, panels a, c). From the peptide and protein detection overlaps alone, one would not expect a large difference in the detected significantly regulated proteins (see ’Results’ section). This suggests that the incomplete spectral library overlap is at the basis of that discrepancy, and the use of different spectral library peptides and associated transitions of peptide ions leads to the difference in quantitative results. At first in contrast to these overlaps, the small overlap of peptides in dataset PXD014943 (Supplementary Figure S5, panel e) is unexpected given that the spectral libraries used in both original and reanalysis were the same (Supplementary Figure S5, panel b). However, upon closer inspection, this could be explained by the fact that the originally reported peptides could have been filtered to a one-peptide-per-protein representation (in the original publication data), as the detected proteins reported showed a larger overlap (Supplementary Figure S5, panel h). Additionally, the dataset PXD014943 showed one of the bigger overlaps in the list of differentially regulated proteins (Supplementary Table S1). This leads to the conclusion that the spectral library composition plays a major role, but other parameters of the analysis also are important for protein detection, and further downstream in protein quantification. In fact, we can highlight one parameter in particular, that has shown a very strong impact on protein quantification values, the method of protein inference (Supplementary Figure S4, Supplementary Tables S3, S2). We chose the ’top3’ method in MSstats, which increased the number of confidently detected proteins and regulated proteins. We would like to strongly encourage future studies to include details and discussion on the method of protein inference to better guide the understanding of presented results to the community. It should also be noted that other differences in methodology, software, and representation of the experimental design of the original study, as detailed in the Methods section, could also be the reason behind the result deviations.

Here we wanted to study feasibility of DIA-MS (re-)analysis with a generic spectral library on a broad range of human datasets. However, the analysis stability is dependent on the use of same or similar spectral libraries (in addition to the acquisition parameters). As shown in this study, and as it should be expected, quantification will vary on the same dataset using different spectral library peptides and associated transitions of peptide ions. The custom (expert) choice of certain peptide ions to include can substantially change the quantitative values observed. For consistent reanalysis pipelines it would be undesirable to choose a custom spectral library for every case, and a stable set of peptides and associated transitions of peptide ions are required for robust results. Our reanalysis results have been made available via Expression Atlas, thereby offering a convenient route to integrate DDA and DIA proteomics expression data with transcriptomics data in the same resource. We hope this will help popularise proteomics approaches in general and DIA approaches in particular, especially for non-proteomics researchers. Finally, the integration of qualitative and quantitative data from complementary *omics* techniques is, in our view, vital for having a broadened understanding of the underlying biological processes in different organisms.

## Methods

### Selection of public DIA proteomics datasets and manual curation

After performing an initial selection of publicly available SWATH-MS datasets from PRIDE^3^ (all measured from human samples with SCIEX instruments), 10 datasets were selected with a preference for studies with technical replicates. Further criteria for reanalysis inclusion were the completeness of data in terms of the necessary files and data types, including both .*wiff* and .*scan* files, existing annotation of the used iRT peptides, and further MS run information, to be able to reconstruct the studies’ experimental design. In all cases, each dataset was manually curated to extract the main analysis characteristics, the processing parameters, the experimental design and sample characteristics. The biological metadata for each dataset was captured in a SDRF file. Then, the raw data files from the selected 10 datasets (Table 1) were downloaded and used as input for the reanalysis (Figure 1a). The main characteristics of the selected 10 datasets are summarised below.

### HCCpct (PXD004873)

This dataset consisted of 76 PCT-SWATH runs from 19 HCC patients’ hepatectomies^41^. In the original analysis, overall 38 tissue samples (pairs of benign and tumorous tissues) were prepared, spiked with iRT peptides and measured with a 5600 TripleTOF instrument over a 45 min LC gradient, in technical replicates. The CAL library was used with the iPortal^42^ software, resulting in 2,579 quantified proteins.

### DigitalKidney (PXD000672)

This kidney dataset consisted of 48 SWATH-MS runs. In the original analysis, 18 kidney tissue samples were processed as benign and tumorous pairs coming from 9 patients with renal cell carcinoma (RCC), six of which were classified as clear-cell RCC (ccRCC), two were classified as papillary RCC (pRCC) and one as chromophobe RCC^16^. Additionally, 4 digests of human kidney tissue were processed in triplicate (twelve aliquots). The processing included spiking with iRT peptides. Data acquisition was performed on a 5600 TripleTOF mass spectrometer over a 120 min LC gradient. The custom target library used contained targets from 41,542 proteotypic peptides, coming from 4,624 proteins, compiled using the TPP^43^ (TransProteomicPipeline) on a DDA analysis of kidney tissues. Using an FDR of 0.1% at a precursor level, 1,632 unique proteins were originally quantified with iPortal.

### BankPrCa (PXD004691)

This dataset contained 224 SWATH-MS runs coming from prostate cancer (PrCa) biopsies^44^. The tissues samples were either fresh frozen (FF) or formalin-fixed-paraffin-embedded before pressure cycling technology (PCT) was used in the original analysis. All samples were spiked with iRT peptides (Biognosys^22^) before injection. Data acquisition was performed on a 5600 TripleTOF mass spectrometer with a 30 min LC gradient, with technical replicates. DIA analysis was performed with a custom spectral library (70,981 peptide precursors from 6,686 UniProtKB/Swiss-Prot proteins) compiled from unfractionated prostate tissue digests (PCT) and measured in DDA mode on the same type of instrument, over a 2 h gradient. The analysis software used was OpenSWATH and iPortal. The original analysis resulted in 3,030 detected proteins and a median CV of 16.2%.

### Wylm (PXD014943)

This PCT-SWATH dataset came from diffuse large B-cell lymphoma (DLBCL)^44^. In the original analysis, the samples were prepared from FFPE and spiked with iRT peptides before the MS data measurement took place. The dataset contained 113 runs, acquired on a 6600 TripleTOF mass spectrometer with a 60 min LC gradient, in technical duplicates. The original data analysis was conducted using a custom target spectral library with iPortal and resulted in 5,769 proteins detected.

### PrCaRegions (PXD003497)

This dataset included 60 PCT-SWATH-MS runs coming from prostatectomies of three individuals^26^. The tissue samples included acinar prostate tumours and a ductal prostate tumour. Prepared samples were spiked with iRT peptides. Data acquisition was performed on a 5600 TripleTOF mass spectrometer with a 120 min LC gradient, with technical replicates for each selected region of the prostatectomies. The target spectral library used for the original analysis was compiled by combining targets from prostate tissue digest runs and the DDA files from the pan-human combined assay library (CAL). In the original analysis, performed with OpenSWATH and TRIC, 6,873 proteins were reported to be consistently quantified, 3,700 proteins of those with a high level of correlation. The dataset included additional SWATH-MS runs of 36 human liver S9 fractions (HLS9) samples and three pooled human liver microsomes (HLM) acquired with the same instrumentation but using a 45 min LC gradient.

### PrCaNetwork (PXD004589)

This dataset contained 210 SWATH-MS runs from PrCa tissue^45^. The tissue material was sourced from 39 individuals. The sample preparation was performed with PCT and spiked with 10% iRT peptides. Data acquisition was performed on a 5600 TripleTOF mass spectrometer, with a 120 min LC gradient with technical replicates. The analysis target library was custom built from 422 DDA MS runs of prostate tissue samples. The original data analysis was performed with OpenSWATH and TRIC, and overall 2,371 quantified proteins were reported.

### BraCaOFLM4 (PXD014194)

This dataset contained 52 breast cancer (BrCa) samples, consisting of 13 estrogen receptor and/or progesterone receptor (ERPR) positive cases, 13 Her2 (Her) positive, 13 estrogen/progesterone/her2 (ERPRHer) positive, and 13 triple negative (TN) breast tumours, and a cohort of 20 ductal carcinoma in situ (DCIS) samples^46^. Overall, 145 SWATH-MS runs were prepared from FFPE samples, measured on a 5600+ TripleTOF mass spectrometer, using a 95 min LC gradient. A custom spectral library containing 3,200 proteins was used in the original analysis performed with Spectronaut PulsarX12 software (Biognosys, Schlieren, Aargau, Switzerland), and resulted in 1,313 detected proteins, of which 1,064 were consistently quantified.

### NCI-60 (PXD003539)

This dataset titled ’Quantitative Proteome Landscape of the NCI-60 Cancer Cell Lines’ consisted of a cell line panel that contained 60 cell lines coming from 9 different tissue types^47^. Each cell line was measured in technical duplicates. The software used for the original analysis was OpenSWATH, with MAYU^48^ and DIA-expert^49^. The 120 MS runs were acquired with the PCT-SWATH method on a 5600 TripleTOF mass spectrometer with a 120 min LC gradient. A custom target spectral library from DDA MS runs obtained from the same samples (containing 86,209 proteotypic peptide precursors in 8,056 proteotypic UniProtKB/Swiss-Prot proteins) was created, resulting in 3,171 consistently detected proteins at a <1% peptide and protein FDR level (with all proteins detected in technical replicates).

### Plasma (PXD001064)

Aiming to use a dataset with a more complex expected variation, a twin study design dataset including longitudinal factors, was also selected^50^. It contained 240 SWATH-MS runs measured on a 5600+ TripleTOF mass spectrometer with a 120 min LC gradient. Of these, 234 were measured from blood sample pairs of 72 monozygotic and 44 dizygotic twins, ranging from 38 to 74 years of age. An additional 6 MS runs were measured from pooled samples as technical replicates. The sampling was done at two different time points: 5.2 ± 1.4 years apart. Originally, a custom target spectral library was created for the original analysis, containing targets coming from 652 proteins detected in mixed plasma samples using DDA experiments and CAL, resulting in targets representing 1,667 unique plasma proteins. An average of 425 proteins were identified using OpenSWATH, 342 of which were consistently quantified in each twin sample at a 1% protein FDR.

### DIATPA (PXD010912)

This human liver dataset consisted of overall 43 SWATH-MS runs^51^. From these 43 runs, 36 were measured from individual human liver S9 fraction (HLS9) samples sourced from 17 males and 19 females, aged 23-81 years, 3 MS runs of pooled HLS9 samples from the same cohort, and 3 MS runs from pooled human liver microsomes (HLM) from 100 males and 100 females, aged 11–83 years. Analysis was done on a 5600+ TripleTOF mass spectrometer with a 90 min LC gradient. A custom target spectral library was constructed from DDA runs of the cohort of 36 samples. At a 1% protein FDR, on average, 1,250 proteins were quantified per each HLS9 sample in the original analysis using Spectronaut Pulsar software (version 11.0, Biognosys).

### SWATH-MS data analysis

The data analysis workflows were constructed using Nextflow. This choice allowed the data processing to be executed either in single-computer mode, on HPC clusters, or on cloud computing platforms. The SWATH-MS data analysis workflow steps (Figure 1b,c) can be broken down into raw data conversion, QC/QA and SWATH-MS window processing, OpenSWATH target generation and data analysis, FDR analysis and multi-run alignment with PyProphet^23^ and TRIC^52^ (Figure 1b), followed by a separate workflow for statistical analysis performed with R and MSstats (Figure 1c), concluded by result inspection and upload to Expression Atlas via custom submission scripts (Figure 1d). The workflows included in the pipeline were built for top-to-bottom analysis, starting with the conversion of the raw instrument data, followed by the statistical data processing with MSstats. The Nextflow workflows were split after the multi-run alignment step and equipped with optional entry points for pre-processed data from OpenSWATH^12^ (.*osw* files) to enable more flexible compute settings. All analysis software was containerised either from available software releases or built-for-purpose to ensure a well-defined compute environment and software compatibility. All container ’recipes’ are included in the GitHub repository (https://github.com/PRIDE-reanalysis/DIA-reanalysis) and can be rebuilt for local use (see Table 3).

**Table 3.** Overview of the containers and software versions used in the open data analysis pipeline.

| Step | Name | URL or DockerHub handle | Version |
| --- | --- | --- | --- |
| Conversion from raw file to mzML | wiffConverter | sciex/wiffconverter:0.7 | 0.7.0 |
| QC/QA | yamato | https://github.com/PaulBrack/Yamato/releases/download/v1.0.4/release-linux-x64.zip | 1.0.4 |
| Window management | python scripts | https://github.com/PRIDE-reanalysis/DIA-reanalysis | 1.0.0 |
| OpenSWATH | OpenSWATH | openswath/Openswath:0.1.2 | 2.4.0 (git 868546e) |
|  | PyProphet |  | 2.0.dev1 (git ddcedac) |
|  | TRIC |  | 0.8.0 (git eeed765) |
| Post-processing | R | https://github.com/PRIDE-reanalysis/DIA-reanalysis | 4.0.3 |
|  | MSstats |  | 3.22.0 |
|  | MyGene.info |  | 1.24.0, Ensembl 99/GRCh38 |

The inputs (Table 4) for the Nextflow pipeline consisted of the SWATH-MS runs as a collection of .*wiff/*.*scan* files, together with a descriptor file in .*TraML* format, detailing the iRT peptides used. As target input, the pipeline consumes a target spectral library file in an OpenSWATH conformant .*tsv* format. In case tool parameters needed changing, an additional .*yaml* file can be provided with updated parameters settings. If a target library in .*pqp* format had already been created or the analysis of processed .*osw* files had already been conducted with different downstream parameters (FDR thresholds), these could be used as alternative entry points in the Nextflow workflow. For the last Nextflow workflow, the study design in an MSstats conformant format must also be provided.

**Table 4.**
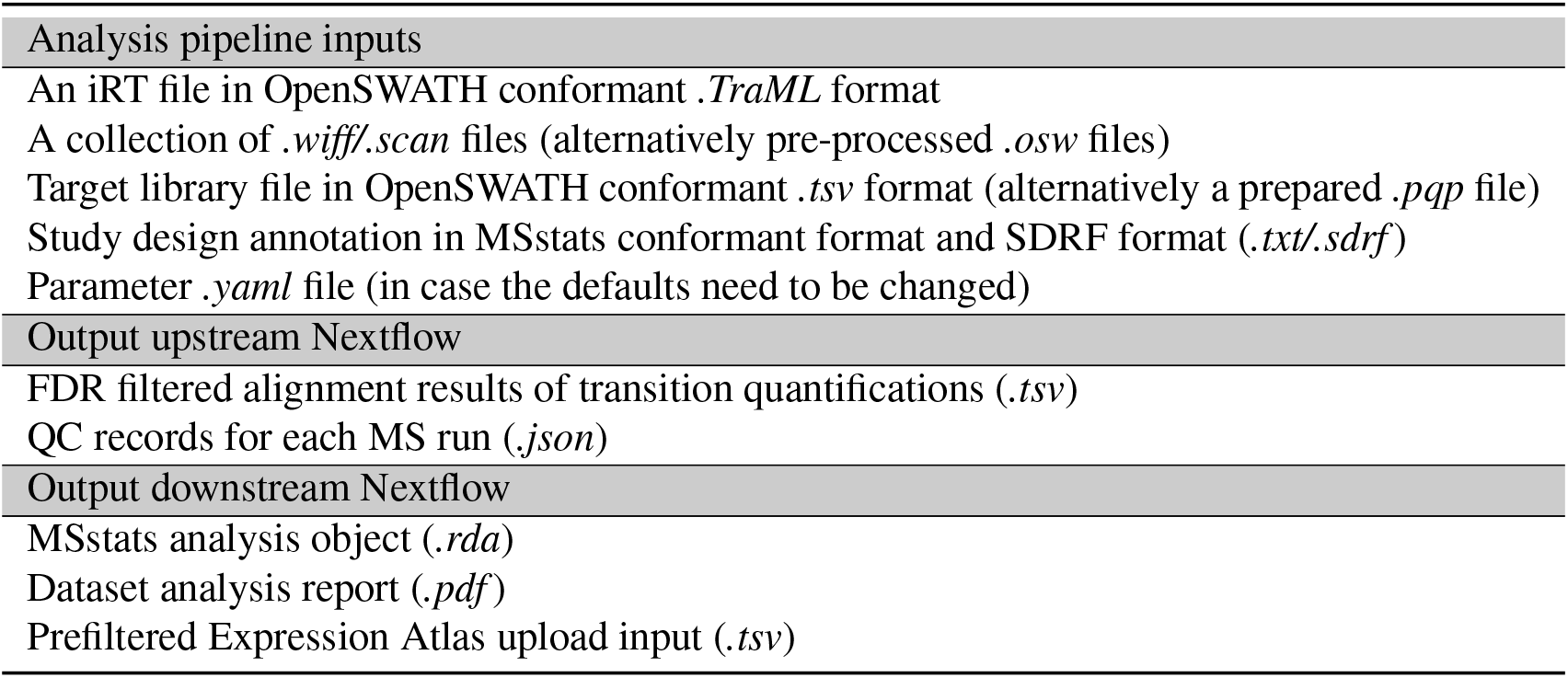
Types of input to the DIA reanalysis pipeline implemented in Nextflow.

### Target spectral library and FDR control and statistical analysis

As target spectral library input for the analysis, we used CAL, which was originally envisioned as a generic large-scale human assay library to support human SWATH-MS studies. It consists of 1,164,312 transitions identifying 139,449 proteotypic peptides and 10,316 proteins, considering only non-redundant entries of UniProtKB/Swiss-Prot. It therefore supports the detection and quantification of 50.9% of all human proteins^15^, as annotated by UniProtKB/Swiss-Prot. It was generated by combining the results from 331 measurements of fractions from different sample types with a technical variation between replicates of below 20% for the quantified signals at a precursor level. For the ’Plasma’ dataset, CAL was filtered for blood/plasma proteins (as annotated in UniProtKB/Swiss-Prot) to reflect the changed base assumption of potentially present proteins. Protein identification was performed with OpenSWATH and PyProphet as described in [^23,53^]. PyProphet was used to calculate the confidence scores on peptide level (global context) and protein level (global context) in parallel at the given FDR thresholds. Decoys were generated with the OpenSWATH tool OpenSwathDecoyGenerator (using default parameters) and the target library assay was customised to the input dataset’s SWATH-MS window setting with OpenSWATHAssayGenerator.

The proteomic analyses were performed at distinct levels (5%, 1% and 0.1%) of peptide and protein FDR, and TRIC was used to align the SWATH-MS runs detected features. The results were then post-processed with MSstats for protein inference and quantitative analysis. As protein inference method, the ’top3’ method was chosen (for the impact of choosing the default ’all’ protein inference method, see the ’Results’ section and Supplementary Figure S4). The abundance values were median normalised and *log*_2_ transformed. Normalisation using MSstats methodology was replaced by a more conservative median normalisation approach. Furthermore, a consistency filter was also applied to filter out proteins with ≤ 50% of all target features of the protein detected over the respective study groups’ MS runs. If the study included multiple study groups and protein expression contrasts of biological interest, additionally, a differential expression analysis was performed, and fold-change contrasts were calculated (see Table 1, ’Analysis type’ column) between study groups. Between the respective groups, the protein *log*_2_ fold-change was computed from the mean of each group’s expression value and the significance level controlled (*α* = 0.05, R software, Welch t-test, Benjamini & Hochberg).

### Technical validation

The resulting protein abundances amongst technical replicates were used to calculate the CV for the datasets. The median CV was calculated per study group and subject, from the median CV for protein abundances in each pair of technical replications and compared to the initially reported CV (Figure 3). For each reanalysed dataset containing technical replicates, the correlation of each study’s pairs of technical replicates was also investigated (Figure 4). Where available, we additionally analysed the correlation of originally reported protein abundances against our reanalysis results. The correlations were measured with the Pearson product-moment correlation coefficient.

### Integration of the results in Expression Atlas

The data integration into Expression Atlas was performed on the results filtered with an FDR threshold of 1% from global context peptide and protein FDR. The MyGene.info R client (version 1.24.0, Ensembl 99/GRCh38) was used to map UniProtKB/Swiss-Prot protein accessions to Ensembl gene IDs. Protein groups with mappings to more than one Ensembl gene ID or decoys and targets without mappings were removed. The filtered and median normalised quantitative results per MS run were integrated into Expression Atlas. In the case of differential datasets, the *log*_2_ fold-changes with *−log*_10_ p-values (adjusted) were also submitted with information on the compared sample groups instead. The corresponding annotated SDRF files are also available. Expression Atlas URLs for each dataset are indicated in Table 1.

## Data availability

All input data was downloaded from PRIDE. Experimental design annotations (.*sdrf*, .*txt*) and reanalysis outputs are available in the reanalysis repository under the respective *PXD* folder. Table 1 contains the collected information on the datasets, Table 5 information on the result availability, Supplementary Table S4 intermediate result availability.

**Table 5.** Detailed information on the datasets included in the reanalysis. *The current URLs correspond to the development instance of Expression Atlas. In the next Expression Atlas release, these datasets will be moved to the production instance. At that point links will be updated to the corresponding URLs starting by https://www.ebi.ac.uk/gxa/experiments/

| Dataset identifier | Originally reported protein quantification | Condition contrasts available | Condition common Proteins | Expression Atlas URL* |
| --- | --- | --- | --- | --- |
| PXD004873 | Per MS run | - | - | https://wwwdev.ebi.ac.uk/gxa/experiments/E-PROT-69/Results |
| PXD000672 | Per MS run | 2 | 975; 1022 | https://wwwdev.ebi.ac.uk/gxa/experiments/E-PROT-59/Results |
| PXD004691 | Per sample | 2 | 1435; 991 | https://wwwdev.ebi.ac.uk/gxa/experiments/E-PROT-68/Results |
| PXD014943 | Per sample | 2 | 3019; 1080 | https://wwwdev.ebi.ac.uk/gxa/experiments/E-PROT-67/Results |
| PXD003497 | Per MS run | - | - | https://wwwdev.ebi.ac.uk/gxa/experiments/E-PROT-66/Results |
| PXD004589 | Per MS run | - | - | https://wwwdev.ebi.ac.uk/gxa/experiments/E-PROT-70/Results |
| PXD014194 | Per MS run | - | - | https://wwwdev.ebi.ac.uk/gxa/experiments/E-PROT-72/Results |
| PXD003539 | - | - | - | https://wwwdev.ebi.ac.uk/gxa/experiments/E-PROT-73/Results |
| PXD001064 | - | - | - | https://wwwdev.ebi.ac.uk/gxa/experiments/E-PROT-60/Results |
| PXD010912 | - | - | - | https://wwwdev.ebi.ac.uk/gxa/experiments/E-PROT-71/Results |

## Code availability

The complete open reanalysis pipeline description and documentation, workflows, container recipes, and custom code and visualisation scripts, as well as parameter input files are available through the GitHub repository at https://github.com/PRIDE-reanalysis/DIA-reanalysis.

## Acknowledgements

We would like to thank all data submitters who made their datasets available in PRIDE. This work has been funded by the BBSRC grant ’Proteomics-DIA’ [grant numbers BB/P024599/1 and BB/P024424/1]. The authors would also like to thank H. Roest for his help in the implementation of the analysis pipeline. JAV, AP and DGS would also like to acknowledge The Wellcome Trust [grant number 208391/Z/17/Z], the EU H2020 grant ’EPIC-XS’ [grant number 823839] and EMBL-core funding.

## Author contributions statement

M.W., A.P., and D.G.S. designed the analysis pipeline, A.P., D.G.S. and M.W. analysed the datasets and integrated the results into Expression Atlas. P.B, P.C., R.L.G. S.J.H, and J.A.V. consulted on the pipeline development, N.G., S.M., P.M., and I.P. contributed to the integration of the data in Expression Atlas. S.J.H, R.L.G. and J.A.V. acquired the funding. M.W, S.J.H., and wrote the manuscript. All authors reviewed the manuscript.

