## Supplementary Material for "Implementing the reuse of public DIA proteomics datasets: from the PRIDE database to Expression Atlas"

Table of contents:


| | |
|---|---|
| Supplementary Table S1 | p. 2 |
| Supplementary Table S2 | p. 3 |
| Supplementary Table S3 | p. 4 |
| Supplementary Table S4 | p. 5 |
| Supplementary Figure S1 | p. 6 |
| Supplementary Figure S2 | p. 7 |
| Supplementary Figure S3 | p. 8 |
| Supplementary Figure S4 | p. 9 |
| Supplementary Figure S5 | p. 10 |

**Supplementary Table S1.** Common protein accessions in significantly differentially expressed proteins in the original analysis and the reanalysis

| PXD | Contrast | Protein Accessions |
|---|---|---|
| PXD000672 | Normal vs ccRCC | P07148, P63104, P42126, P10809, P48047, Q86YH6, Q92597, P00740, P50238, P99999, P09110, Q9UHK6, P24539, Q02252, P43487, P22352, P08195, O75947, P04406, P45954, P47914, Q16718, Q16775, Q03154, O43175, Q9ULZ3, P40121, P24752, Q9H0W9, P18669, P25705, Q5T2W1, O75964, P00558, P15924, O95831, Q6PCB0, P07237, P09467, P01023, P30038, Q9H299, P20711, P06748, Q00325, P08758, P21912, P31937, P36543, P02792, P46777, P00966, P15880, P24821, Q13825, P06576, O75340, Q04941, P00918 |
| PXD000672 | ccRCC vs pRCC | Q99623, P00367, P35232, P55084, Q16891, P40939, P30044, Q9Y277, Q9UJZ1, P10253, Q13642, P53597, Q08380, P09110, Q13011, P10809, Q5VWZ2, P61604, P54819, P38117, P02794, P56385, O75390, O75964, Q16836, P31040, Q07021, Q9H2U2, P49748, P30084, P30049, P30048, Q16698, P36542, P40926, Q02252, Q9HCC0, P48047, P05091, Q00325, P12277, O43615, P06576, P05141, Q6IAA8, P11310, Q86SX6, Q99798, O95831, P25705, P36957, O75947, P49411, Q9Y2X3, P11177, P00403, Q9BX68, P10606, P13073, P99999, P42126, P18859, P24752, P24539, Q8NE62, P00505, P14406, Q9Y5L4, P45954, Q7Z4W1, Q13564, Q6P587, P08574, Q13423, P00846, P12235, P50440, P82933, O43169, Q9UHK6, Q969I3, P09669, O14880, Q13825, P12110, Q14315, P19827, P02656, Q15121, P10412, P00738, P50454, Q92597, P51884, P67936, O60701, P05155, P60981, P00338, P20962, Q15847, Q9H299, P02790, P01876, P30101, P35579, Q93088, P60660, P00558, P01024, O00299, P09525, P09493, P01834, P14618 |
| PXD014943 | eDLBCL vs PCNSL | P43403, P13760, Q15582, P14136, P04271, Q13885, P09543, P12277, Q9BVA1, P09471, P09936, Q12860, Q16555, P29992, Q12765, Q13449, P01871, Q8IXJ6, Q9UM22, P80723, P00568 |
| PXD004691 | Normal vs tumour (fresh frozen) | Q9BUD6, O95994, Q15063 |
| PXD004691 | Normal vs tumour (paraffin embedded) | P06748, P07585, P13647, P15309, P17661, P40926, Q04837, Q15063, Q8NBJ4 |

**Supplementary Table S2.** Quantified protein numbers per study using the ’top3’ protein inference setting

| Dataset | FDR | Reanalysis proteins | Original - before filter | Original - after filter | Reanalysis proteins (‘50% missing) | Reanalysis proteins (‘50% per group) | Peptides | Comment - original data |
|---|---|---|---|---|---|---|---|---|
| PXD004873 | 1% | 3,530 | N/A | 2,579 | 3,530 | 3,392 | 22,231 | 0.1% FDR threshold |
| PXD000672 | 1% | 4,053 | N/A | 1,632 | 4,053 | 4,037 | 31,888 | 0.1% FDR threshold |
| PXD004691 | 1% | 2,872 | N/A | 3,030 | 2,872 | 2,686 | 17,943 | No consistency filter mentioned |
| PXD014943 | 1% | 5,946 | N/A | 5,769 | 5,946 | 5,568 | 45,986 | No consistency filter mentioned |
| PXD003497 | 1% | 2,754 | 6,873 | 3,700 | 2,752 | 2,704 | 20,296 | 6,873 filtered to 3,700 for subsequent analyses |
| PXD004589 | 1% | 3,703 | N/A | 2,371 | 3,702 | 3,298 | 31,727 | Number of proteins mentioned in supplementary material |
| PXD014194 | 1% | 2,239 | 1,313 | 1,064 | 2,238 | 2,145 | 11,593 | 1313 filtered to 1064 for subsequent analyses |
| PXD003539 | 1% | 7,097 | 6,556 | 3,171 | 7,096 | 6,867 | 77,014 | Total number of proteins mentioned in supplementary material only (supplementary figure 2) |
| PXD001064 | 1% | 207 | 425 | 342 | 207 | 197 | 3,508 | 425 on average, 342 selected for consistency |
| PXD010912 | 1% | 4 ,224 | N/A | 1,250 | 4,224 | 4,123 | 34,776 | Manuscript text provides protein number approximation and value range for individual runs |

**Supplementary Table S3.** Quantified proteins per study using the 'all' protein inference setting

| Dataset | FDR | Reanalysis proteins | Original - before filter | Original - after filter | Reanalysis proteins ('50% missing) | Reanalysis proteins ('50% per group) | Comment - original data |
|---|---|---|---|---|---|---|---|
| PXD004873 | 1% | 3,530 | N/A | 2,579 | 2,925 | 2,297 | 0.1% FDR threshold |
| PXD000672 | 1% | 4,053 | N/A | 1,632 | 3,932 | 2,648 | 0.1% FDR threshold |
| PXD004691 | 1% | 2,872 | N/A | 3,030 | 2,033 | 1,688 | No consistency filter mentioned |
| PXD014943 | 1% | 5,946 | N/A | 5,769 | 4,336 | 3,879 | No consistency filter mentioned |
| PXD003497 | 1% | 2,754 | 6,873 | 3,700 | 2,576 | 2,109 | 6,873 filtered to 3,700 for subsequent analyses |
| PXD004589 | 1% | 3,703 | N/A | 2,371 | 2,043 | 2,366 | Number of proteins mentioned in supplementary material |
| PXD014194 | 1% | 2,239 | 1,313 | 1,064 | 1,658 | 1,001 | 1,313 filtered to 1064 for subsequent analyses |
| PXD003539 | 1% | 7,097 | 6,556 | 3,171 | 5,412 | 4,299 | Total number of proteins mentioned in supplementary material only (supplementary figure 2) |
| PXD001064 | 1% | 207 | 425 | 342 | 176 | 174 | 425 on average, 342 selected for consistency |
| PXD010912 | 1% | 4,224 | N/A | 1,250 | 3,915 | 2,924 | Manuscript text provides protein number approximation and value range for individual runs |

**Supplementary Table S4.** Availability and location of intermediate results per study.

| Dataset Identifier | .tsv TRIC output location |
|---|---|
| PXD004873 | https://uk1s3.embassy.ebi.ac.uk/DIA-reanalysis/intermediate-results/tric_tsvs/PXD004873.tric.tsv.tar.gz |
| PXD000672 | https://uk1s3.embassy.ebi.ac.uk/DIA-reanalysis/intermediate-results/tric_tsvs/PXD000672.tric.tsv.tar.gz |
| PXD004691 | https://uk1s3.embassy.ebi.ac.uk/DIA-reanalysis/intermediate-results/tric_tsvs/PXD004691.tric.tsv.tar.gz |
| PXD014943 | https://uk1s3.embassy.ebi.ac.uk/DIA-reanalysis/intermediate-results/tric_tsvs/PXD014943.tric.tsv.tar.gz |
| PXD003497 | https://uk1s3.embassy.ebi.ac.uk/DIA-reanalysis/intermediate-results/tric_tsvs/PXD003497.tric.tsv.tar.gz |
| PXD004589 | https://uk1s3.embassy.ebi.ac.uk/DIA-reanalysis/intermediate-results/tric_tsvs/PXD004589.tric.tsv.tar.gz |
| PXD014194 | https://uk1s3.embassy.ebi.ac.uk/DIA-reanalysis/intermediate-results/tric_tsvs/PXD014194.tric.tsv.tar.gz |
| PXD003539 | https://uk1s3.embassy.ebi.ac.uk/DIA-reanalysis/intermediate-results/tric_tsvs/PXD003539.tric.tsv.tar.gz |
| PXD001064 | https://uk1s3.embassy.ebi.ac.uk/DIA-reanalysis/intermediate-results/tric_tsvs/PXD001064.tric.tsv.tar.gz |
| PXD010912 | https://uk1s3.embassy.ebi.ac.uk/DIA-reanalysis/intermediate-results/tric_tsvs/PXD010912.tric.tsv.tar.gz |

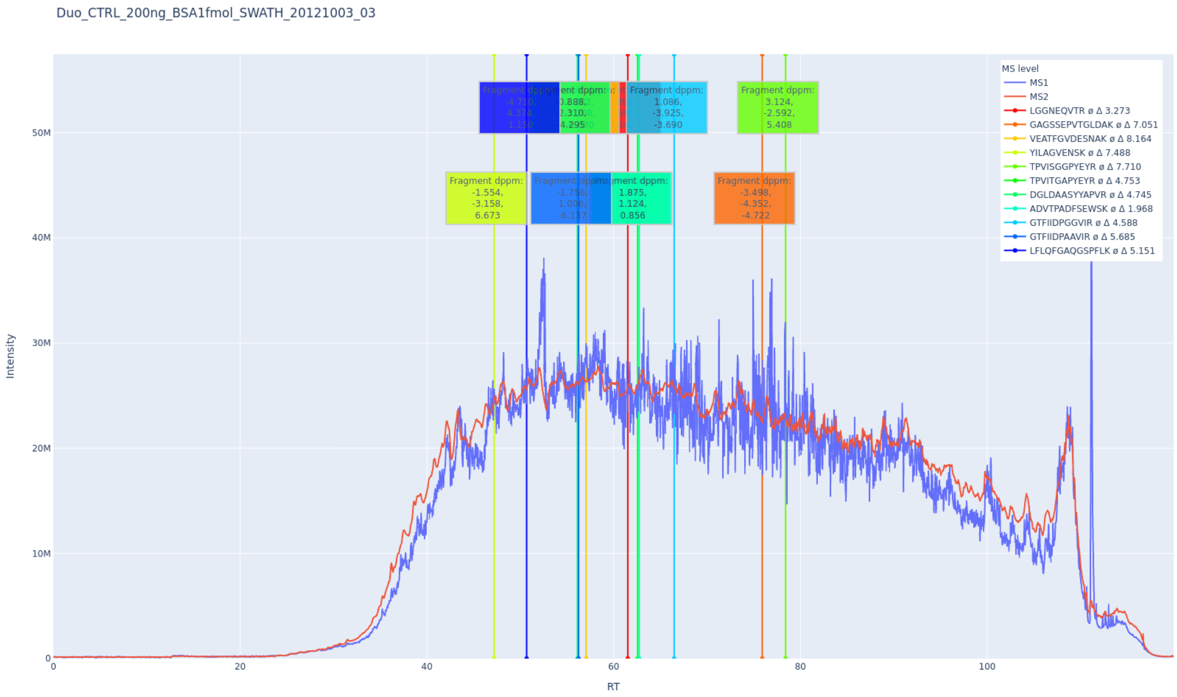


**Supplementary Figure S1.** QC plot of a SWATH-MS run from dataset PXD001506 with containing MS1 and MS2 TIC visualisation and labeled iRT peptide targets. A colour gradient from red to blue indicates the expected order of iRT peptides.

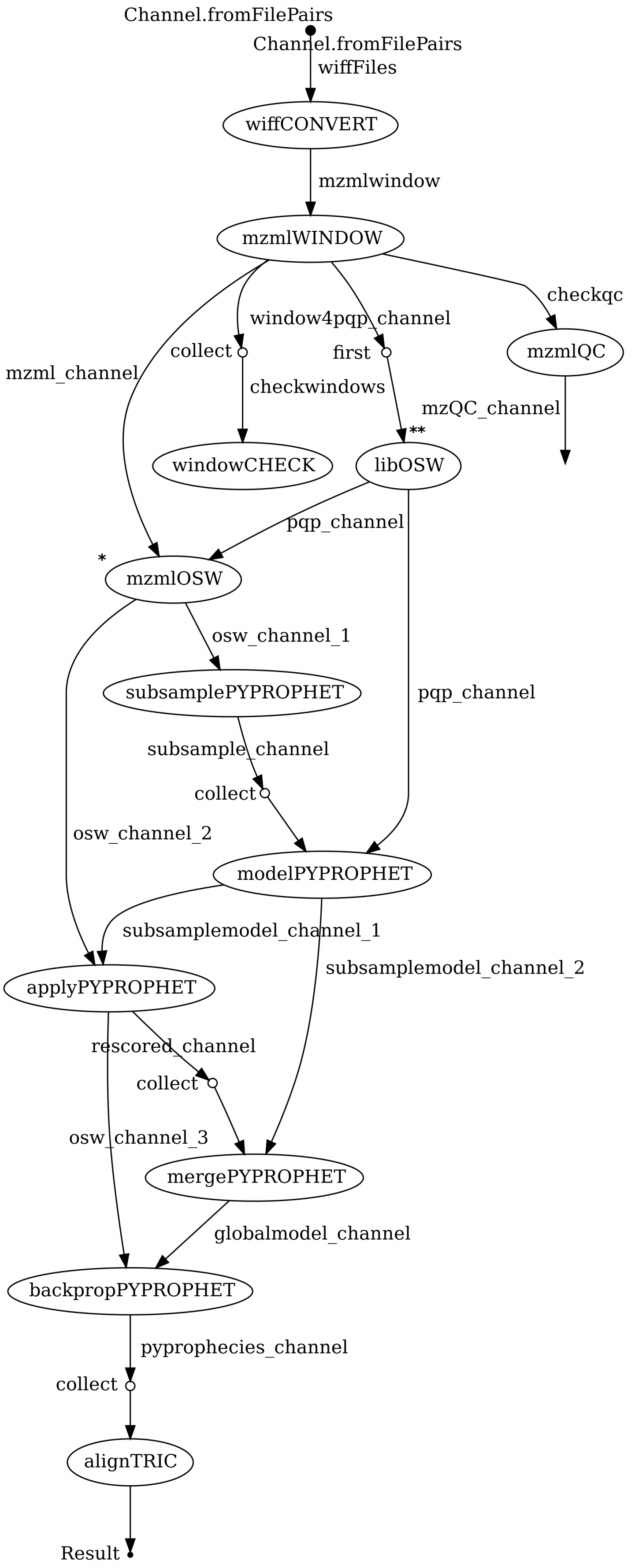


**Supplementary Figure S2.** Detailed visualisation of the upstream Nextflow workflow step sequences. Optional entry points with alternative inputs are marked with an asterisk.

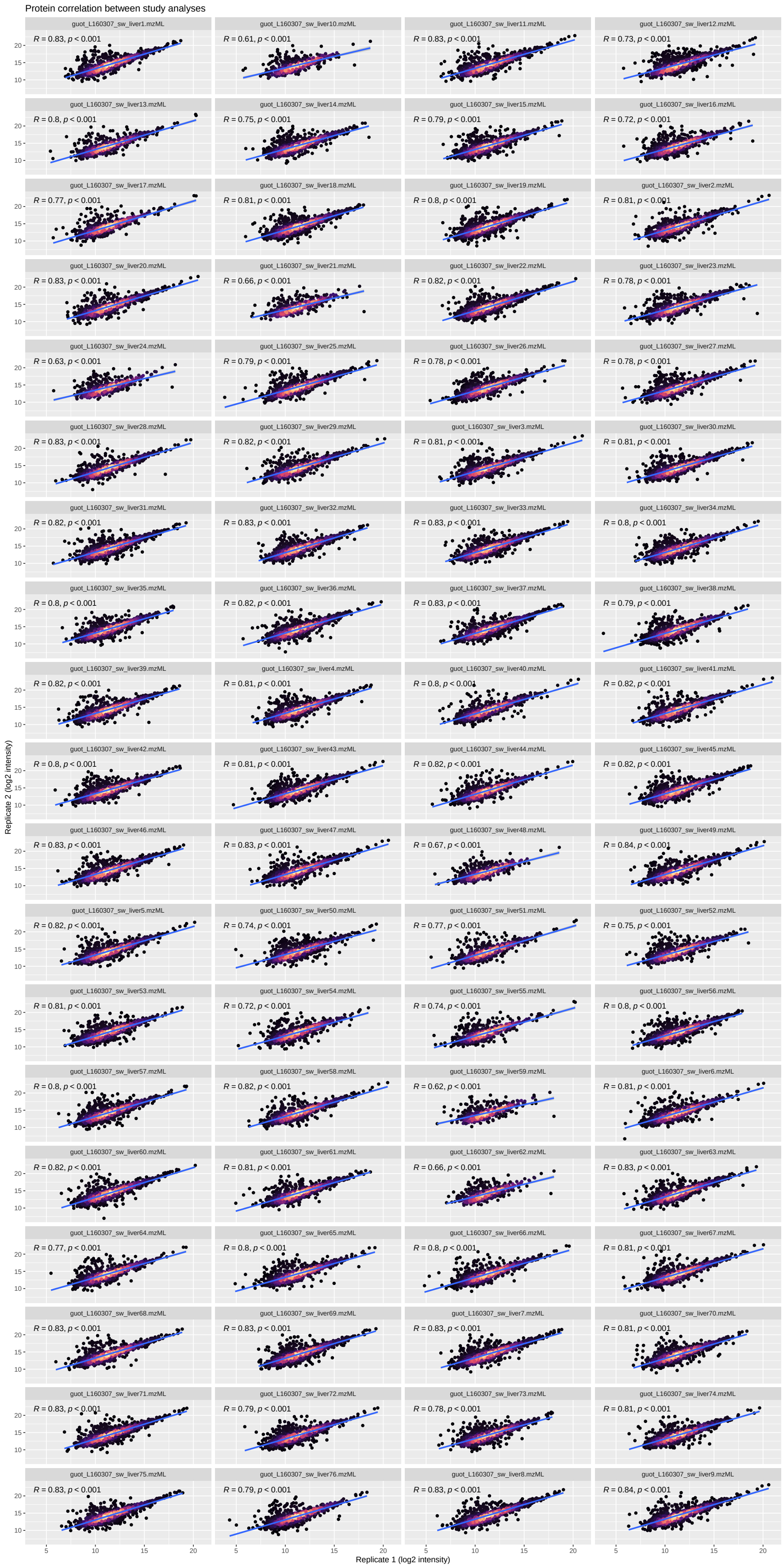

**Supplementary Figure S3.** Run-wise correlation of protein quantification results, reanalysed protein Intensities versus originally published protein intensities. Each one close-to-zero (< 0.05) intensity pair was removed from 5 runs (guot_L160307_sw_liver32.mzML, guot_L160307_sw_liver34.mzML, guot_L160307_sw_liver67.mzML, guot_L160307_sw_liver7.mzML, guot_L160307_sw_liver70.mzML).

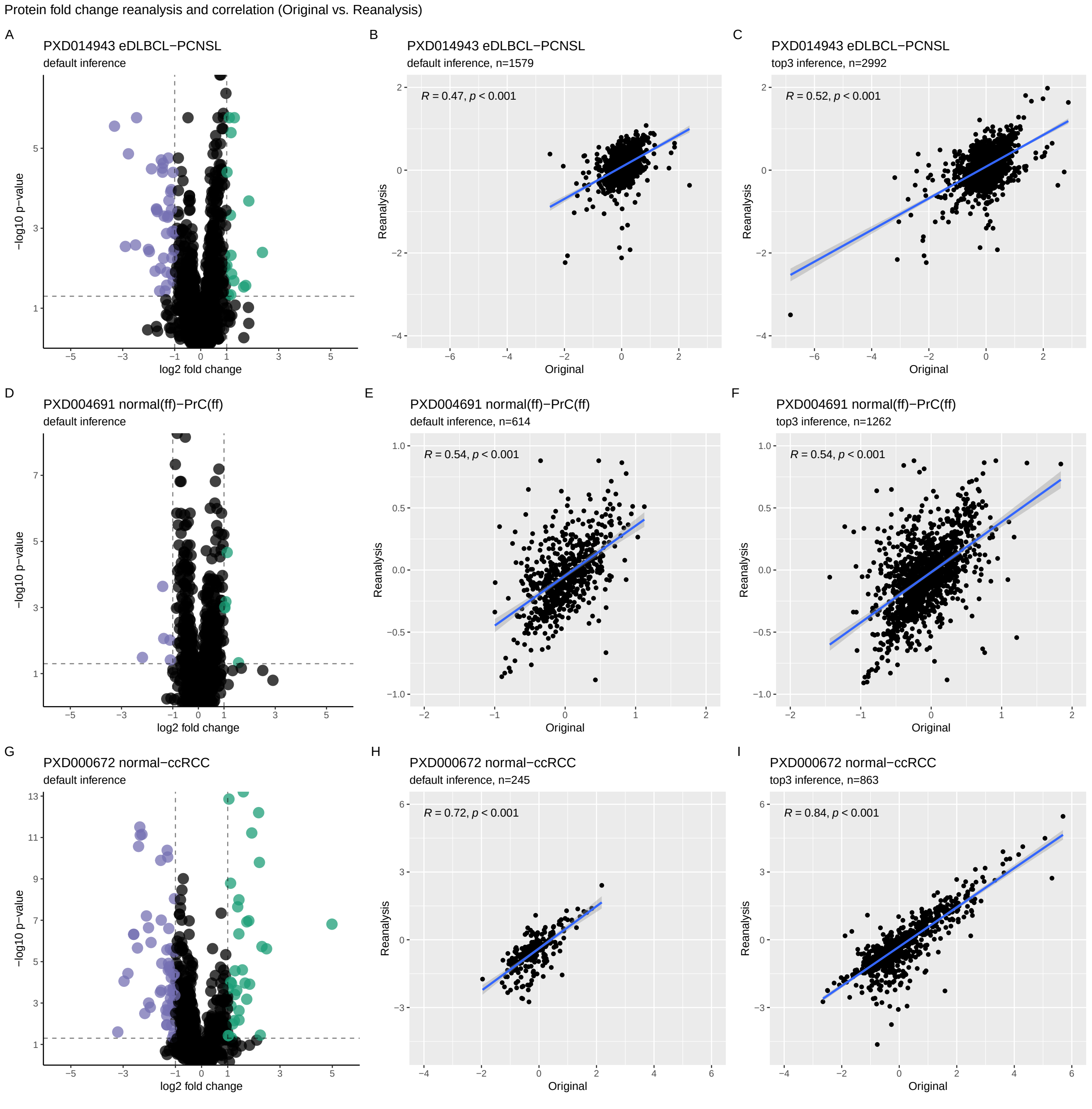


**Supplementary Figure S4.** Inference method influence on differential expression analysis. A, D, G) Volcano plots corresponding to 'all' protein inference method; B, E, H) Fold change (FC) log2 value correlations corresponding to 'all' protein inference method reanalysis vs original; C, F, I) FC log2 value correlations corresponding to 'top3' protein inference method reanalysis vs original. The FC compared are represented by points on the plot. Significant FC proteins are colour indicated. The dashed lines indicate the fold-change cutoff of 2 and the p-value cutoff at 0.05.

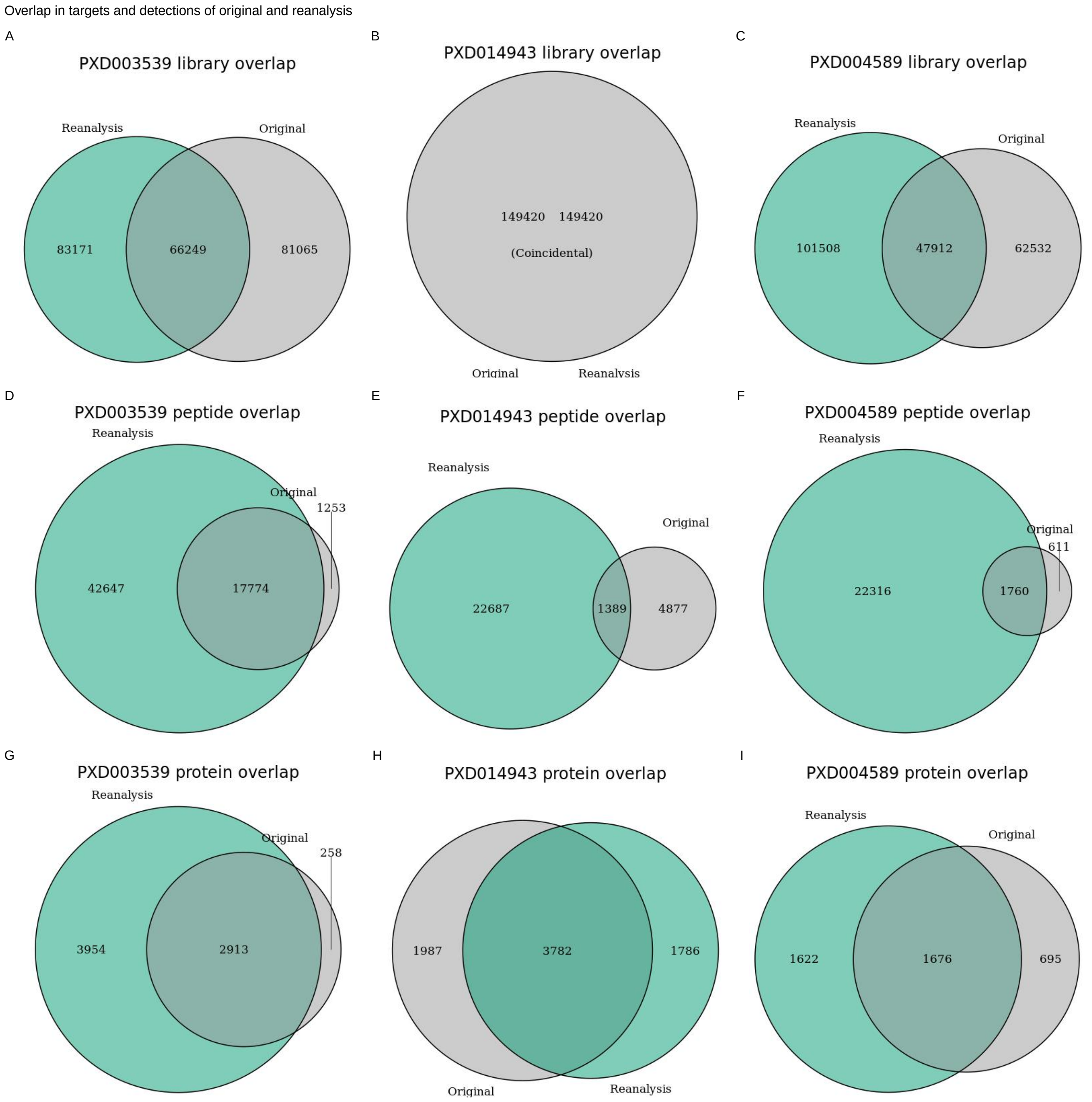


**Supplementary Figure S5.** Target library (A,B,C), peptide detection (D,E,F), and protein detection (G,H,I) overlaps for PXD003539, PXD014943, and PXD004589 between the originally published results and the reanalysis.
